# Measuring individual identity information in animal signals: Overview and performance of available identity metrics

**DOI:** 10.1101/546143

**Authors:** Pavel Linhart, Tomasz Osiejuk, Michal Budka, Martin Šálek, Marek Špinka, Richard Policht, Michaela Syrová, Daniel T. Blumstein

## Abstract

1. Identity signals have been studied for over 50 years but there is no consensus as to how to quantify individuality. While there are a variety of different metrics to quantify individual identity, or individuality, these methods remain un-validated and the relationships between them unclear.
2. We contrasted three univariate and four multivariate metrics (and their different computational variants) and evaluated their performance on simulated and empirical datasets.
3. Of the metrics examined, Beecher’s information statistic (H_S_) was the best one and could easily and reliably be converted into the commonly used discrimination score (and vice versa) after accounting for the number of individuals and calls per individual in a given dataset. Although Beecher’s information statistic is not entirely independent of sampling parameters, this problem can be removed by reducing the number of parameters or by increasing the number of individuals.
4. Because it is easily calculated, has superior performance, can be used to describe single variables or signal as a whole, and because it tells us the maximum number of individuals that can be discriminated given a set of measurements, we recommend that individuality should be quantified using Beecher’s information statistic.

## Introduction

The fact that conspecific individuals differ in consistent ways underlies a number of theoretically important questions in biology such as explaining cooperative behavior or understanding the evolution of sociality (Crowley et al., 1996; Bradbury & Vehrencamp, 1998; Tibbetts, 2004). Because it may be advantageous for animals to choose with whom they interact or respond to (Wilkinson, 1984; Godard, 1991), there may be selection both to produce individually-distinctive signals and to discriminate among them (Tibbetts & Dale, 2007; Wiley, 2013). Individually-distinctive traits can also be used to help wildlife population censuses or to monitor individuals (Terry & McGregor, 2002; Blumstein et al., 2011). For these purposes, identity information in animal signals has been quantified by several different univariate and multivariate metrics, especially in the acoustic domain (Miller, 1978; Hafner, Hamilton, Steiner, Thompson, & Winn, 1979; Beecher, 1989; Searby & Jouventin, 2004; Mathevon, Koralek, Weldele, Glickman, & Theunissen, 2010).

For identity signals to function properly, they should maximize the between-individual variation and minimize the within-individual variation. Therefore, to quantify an individual’s identity we require repeated measurements of one or more traits on a given set of individuals within a population. This is well acknowledged in the study of acoustic signals (e.g., Hutchison, Stevenson, & Thorpe, 1968; Beecher, 1989; Robisson, Aubin, & Bremond, 1993). A typical study of acoustic identity signaling would record large number of vocalizations from each individual under different conditions (different time intervals, distances, etc.), measure a set of acoustic traits (e.g., fundamental frequency, duration, formant structure, frequency modulation, etc.), and then calculate the individual identity either directly through comparing between and within individual variation, or indirectly through discrimination between individuals. In studies of chemical or visual signals, robust assessment of within-individual variation by having many replicates from a single individual remains uncommon (Kondo & Izawa, 2014; but see, e.g., Kean, Chadwick, & Müller, 2015) although quantification of individual identity might be expected in future studies.

A variety of identity metrics have proliferated because the existing metrics were considered biased (Beecher, 1989; Mathevon et al., 2010) or unsuitable for a particular signal type (Searby & Jouventin, 2004). Furthermore, different equations have been sometimes used to calculate the same identity metric (Beecher, 1989; Lein, 2008; Charrier, Aubin, & Mathevon, 2010; Linhart & Šálek, 2017). Thus there is no consensus about how to properly measure identity. As a result, researchers have generally avoided quantitative comparisons between studies (Insley, Phillips, & Charrier, 2003), although there have been a few of using exactly the same methods for several different species (Beecher, Medvin, Stoddard, & Loesche, 1986; Lengagne, Lauga, & Jouventin, 1997; Pollard & Blumstein, 2011). The lack of a commonly used identity metric is a major impediment toward understanding the evolution of identity signaling and indeed, the evolution of individuality.

Here we review previously developed univariate and multivariate metrics that have been used to quantify individual identity information in signals and we test their performance on simulated and empirical datasets. In particular, we investigated the following metrics: F-value, Potential of individual coding PIC, Beecher’s information statistic H_S_, Efficiency of modulated signature H_M_, and Mutual information MI. We further evaluated different computational variants found in literature in case of PIC and H_S_ (see Methods and Supplement 1 for a detail overview of metrics and their variants).

We compare the performance of metrics to a hypothetical ideal identity information metric. We propose that ideal identity metric should have two basic characteristics: 1) it should not be systematically biased by study design (no systematic effects of number of individuals in a study and number of calls per individual in a study); and 2) in the multivariate case (i.e., when it is used to quantify individuality based on measurements of multiple signal features), it should rise with number of meaningful parameters and decrease with covariance between them. Also, for both univariate and multivariate case, we expect the metric will have a meaningful zero in case there is no identity content in a signal. Finally, we expect no upper limit on the degree of individuality; in theory, and given sufficient variation and variables, one could discriminate among an infinite number of individuals. We also wished to see if each of two commonly used metrics (Beecher’s information statistic H_S_, and discrimination score DS) could be converted to the other metric to facilitate comparative analyses of the evolution of individuality.

## Material and methods

We used R for simulations and statistical analysis (R Core Team, 2012). Our simulated and empirical data along with analysis scripts are available on GitHub (Linhart, 2018).

### Datasets

#### Simulated datasets

We constructed datasets with univariate and multivariate normal distributions with parameters covering wide range of values – individuality (id = 0.01, 1, 2.5, 5, 10), number of observations / calls per individual (o = 4, 8, 12, 16, 20), number of individuals (i = 5, 10, 15, 20, 25, 30, 35, 40), and, for multivariate datasets, the covariance among variables (cov = 0, 0.25, 0.5, 0.75, 1) and the number of variables (p = 2, 4, 6, 8, 10). Individuality (id) represents ratio of standard deviations between and within individuals (id = SD_between_ / SD_within_; SD_between_ was calculated from means for each individual). A single covariance (cov) value was used in the variance-covariance matrix to define covariances between all pairs of variables (detailed description in Supplement 2). We asked how dataset parameters (i, o, p, cov, id) influenced the value of each identity metric. To explore this, all combinations of dataset parameters were exhaustively sampled with 20 iterations on each unique combination of parameters. In each iteration, a new dataset was generated to ensure independence between samples. We developed R scripts involving “rnorm” and MASS package (Venables & Ripley, 2002) “mvrnorm” function to generate the datasets.

**Figure 1.**
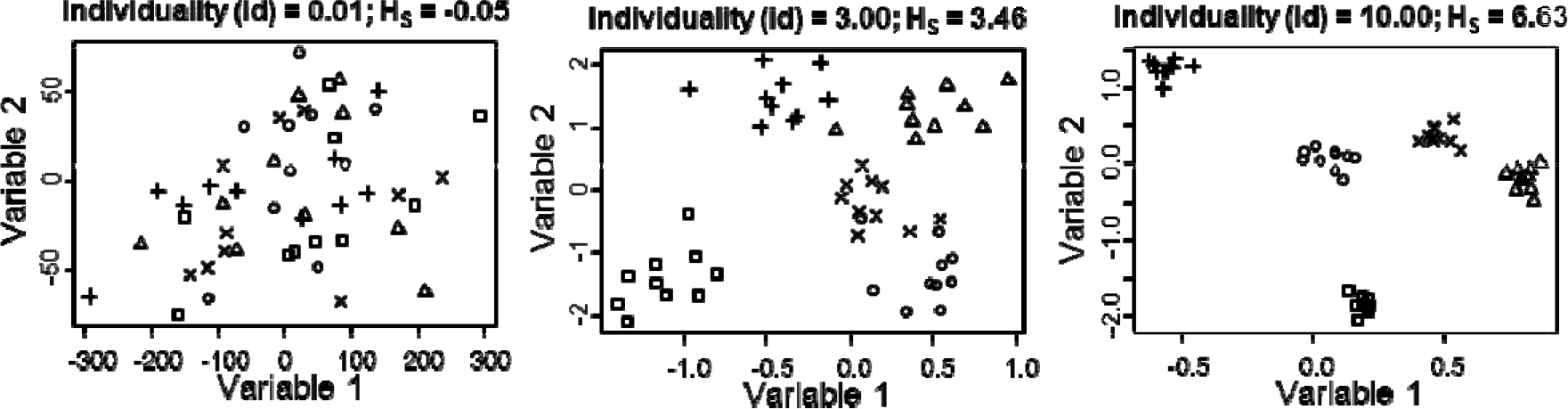
Illustration of three artificial multivariate datasets that differ only in the individuality used to generate datasets. Settings for the function generating these datasets: i = 5, o = 10, p = 2, cov = 0, id = 0.01, 3, and 10

#### Empirical datasets

We used six datasets from four different species: little owls *Athene noctua* (ANmodulation, ANspec) (Linhart & Šálek, 2017), corncrake *Crex crex* (CCformants, CCspec) (Budka & Osiejuk, 2013), yellow-breasted boubous *Laniarius atroflavus* (LAhighweewoo) (Osiejuk et al. unpublished data), and domestic pigs *Sus scrofa* (SSgrunts) (Syrová, Policht, Linhart, & Špinka, 2017) (Figure 2). In two species – corncrakes and little owls – calls were described by two different sets of variables. In little owls, we described calls by frequency modulation (ANmodulation) or parameters describing the distribution of the frequency spectrum (ANspec). In corncrakes, we used formants (CCformants) and parameters describing the distribution of the frequency spectrum (CCspec). Because datasets varied with respect to the number of individuals (33 – 100) and the number of calls per individual available (10 – 20), we scaled all datasets down to lowest common denominator by randomly selecting individuals and calls from bigger datasets. Eventually, each dataset had 33 individuals and 10 calls per individual. Each dataset also used different numbers of variables to describe the calls’ acoustic structure (ANmodulation = 11, ANspec = 7, CCformants = 4, CCspec = 7, LAhighweewoo = 7, SS grunty = 10). In all these empirical datasets, assumptions of multivariate normality were tested (Korkmaz, Goksuluk, & Zararsiz, 2014), but not met. This issue is common for research studies on acoustic individual identity. Authors deal with it by eliminating problematic variables (e.g., Sousa-Lima, Paglia, & da Fonseca, 2008; Couchoux & Dabelsteen, 2015), using non-parametric classification methods (e.g., Tripovich, Rogers, Canfield, & Arnould, 2006; Mielke & Zuberbuehler, 2013), or by relying on robustness of cross-validated DFA towards relaxed assumptions (e.g., Mathevon et al., 2010; Schneiderová, 2012). We used the last approach. If the assumptions of discriminant analysis are not met the results should be less stable when using different sampling and hence our results should be conservative.

**Figure 2.**
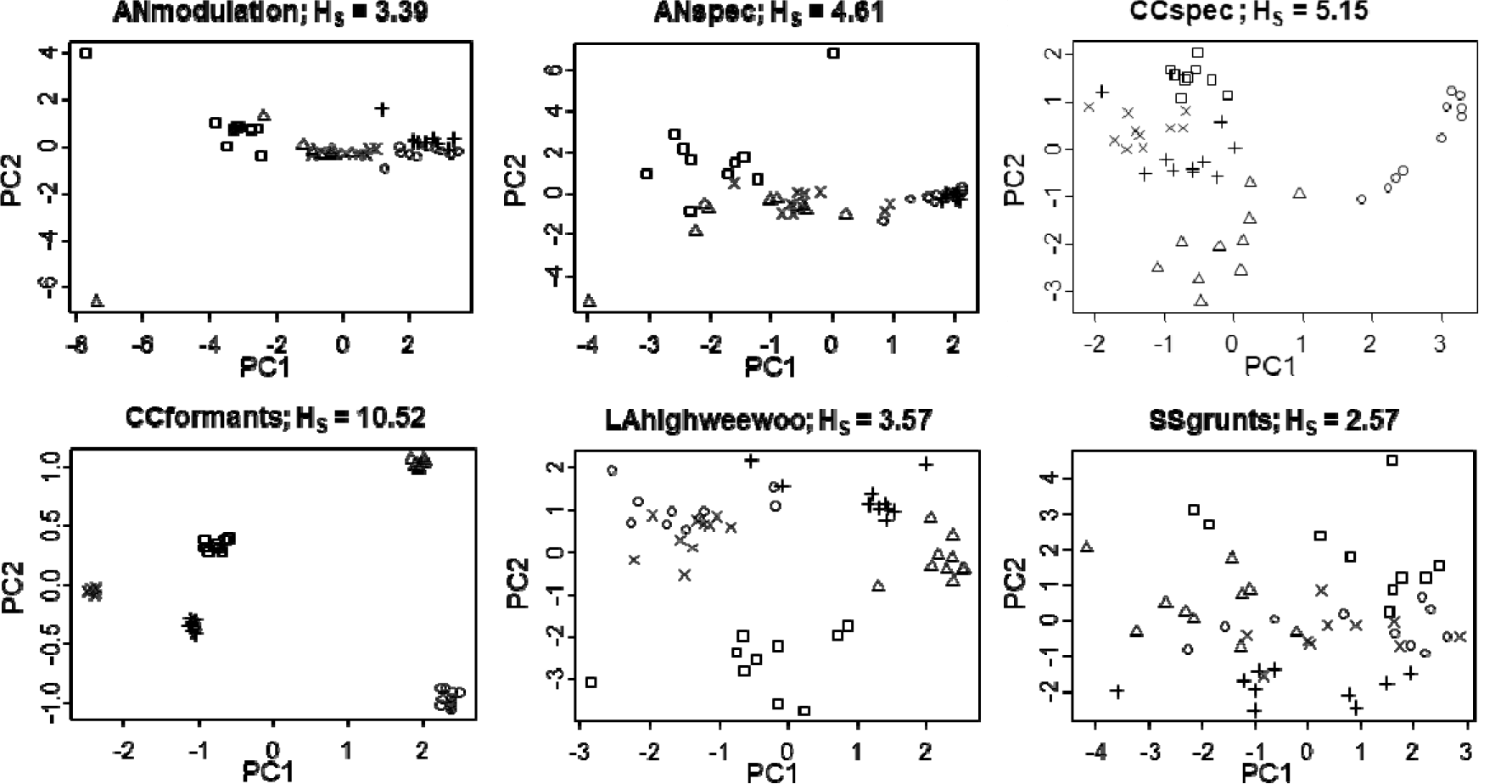
Illustration of empirical datasets. Five individuals were randomly sampled from each dataset of 33 individuals and all 10 calls per individual were selected. H_S_ for a full dataset is shown. Data were centered and scaled and subjected to PCA. The first two Principal Components are plotted.

### R functions to calculate individuality metrics

The following scripts were used to calculate seven variants of three univariate metrics: F value (calcF), Potential of individual coding PIC (calcPICbetweentot, calcPICbetweenmeans), and Beecher’s information statistic (calcHSntot, calcHSnpergroup, calcHSngroups, calcHSvarcomp). PIC is defined as a ratio of between-individual to within-individual coefficients of variation (e.g., Robisson et al., 1993; Lengagne et al., 1997):

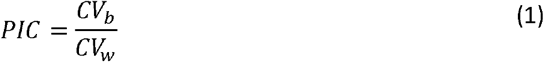

Two variants of PIC differ in whether CV_b_ in the formula is calculated from all values (PIC_betweentot_) (e.g., Charrier et al., 2010), or means for each individual are calculated first and CV_b_ is then calculated from these means (PIC_betweenmeans_) (e.g., Lein, 2008). H_S_ is based on F-value but unlike F-value, H_S_ accounts for sample size:

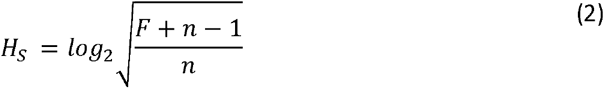

The source of confusion is the ‘n’ in the formula. Total sample size (H_Sntot_), number of groups (i.e., individuals) (H_Sngroups_), and number of samples per group (H_Snpergroup_) could all be used as ‘n’ in this equation. Some studies explicitly state they used number of individuals as ‘n’ (e.g., Pollard, Blumstein, & Griffin, 2010; Linhart & Šálek, 2017), but the properties of H_S_ values in these studies did not match the properties suggested in the original article by Beecher (1989). Yet another approach to calculate H_S_ is to extract the variance component estimates and use the total (⍰_T_) and the residual variance (⍰_W_, associated with random factor) to calculate H_S_ (H_Svarcomp_) (Beecher, 1989; Carter, Logsdon, Arnold, Menchaca, & Medellin, 2012):

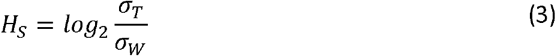

The following scripts were used to calculate multivariate metrics: calcDS, calcHSnpergroup, calcHM, calcMI. The calcDS is based on ‘lda’ (‘MASS’ package). The calcMI function uses ‘lda’ (‘MASS’ package) and ‘mutinformation’ (‘infotheo’ package).

Multivariate identity metrics were always calculated from data (simulated or empirical) that were centered to have a mean of zero, scaled to unit variance, and subjected to principal component analysis.

### Statistical analysis

Our goal was to ask whether there are systematic biases for each identity metric given different parameters that reflect sampling design. The relationship between a given identity metric and each of the parameters was assessed graphically by plotting the mean value and the 95% confidence intervals of an identity metric against all of the modelled data parameters separately. We then used a one-way ANOVA to test whether an identity metric was constant across all levels of a parameter. If we found significant differences, we followed up these with post-hoc Tukey tests to identify which parameter levels differed. Due to high number of comparisons, we only reported comparisons of neighboring parameter levels. We used linear and non-parametric loess regression to convert H_S_ to DS and vice versa. Loess regression included the number of individuals and number of calls per individual as additional predictors. We used Spearman correlation coefficients to quantify between-metric consistency of ranking individuality in datasets. Pearson correlations were used to assess consistency within identity metrics in full and partial datasets. We then used Friedman test, followed by a series of Wilcoxon tests (for post-hoc comparison of differences between levels), to compare correlation coefficients obtained for each pair of the metrics.

## Results

The comparison of available univariate and multivariate metrics to an ideal metric is shown in Table 1.

**Table 1.**
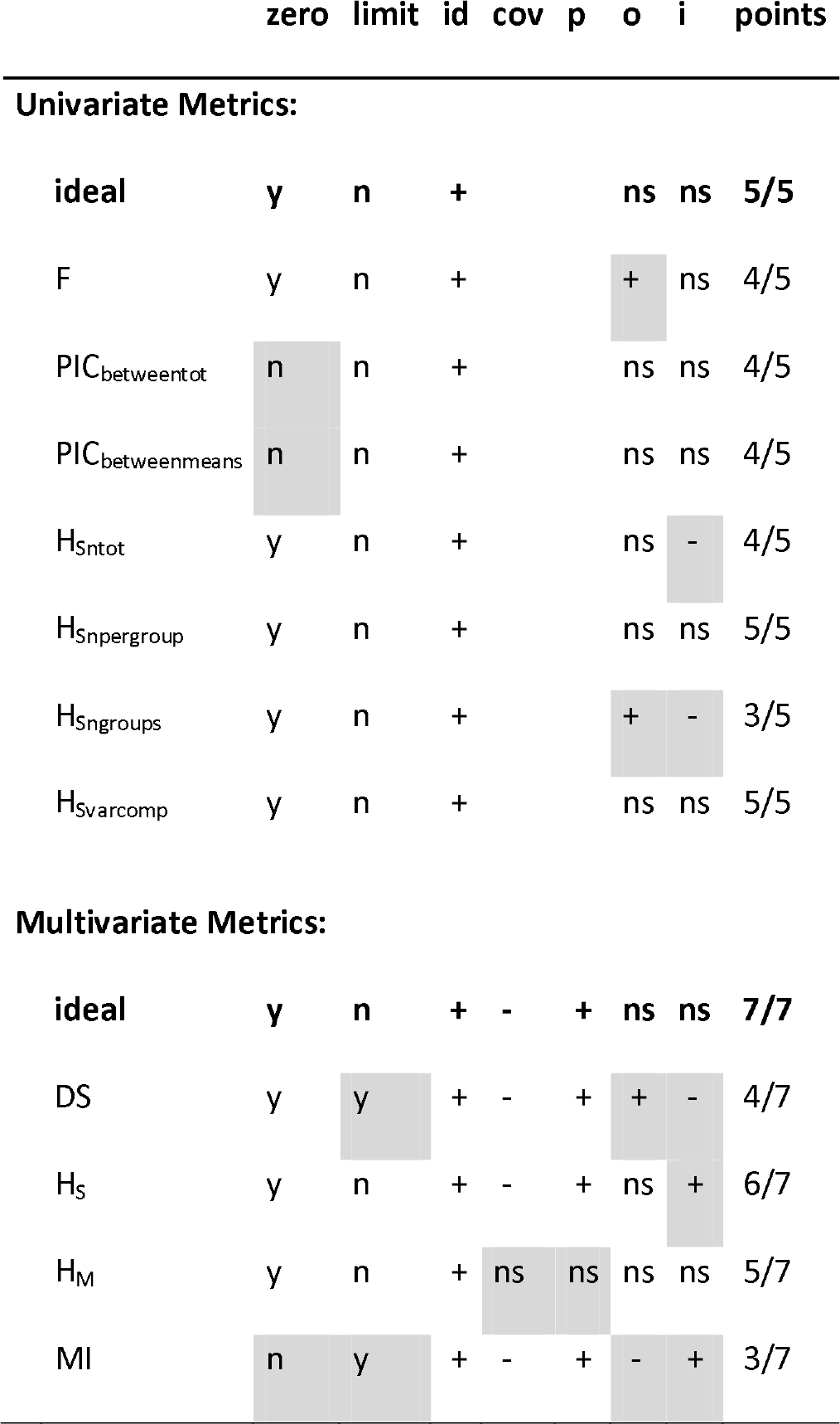
The comparison of available univariate and multivariate metrics to a hypothetical ideal metric. We summed the number of matches (points) to compare different metrics to the ideal metric. Non-matching cells are highlighted in grey background. ‘zero’ – metric has a meaningful zero; ‘limit’ – metric is limited from the top by an asymptote; ‘id’ – change in response to increasing identity information in data; ‘cov’ – response to increasing covariance between variables; ‘p’ – response to increasing number of variables; ‘o’ – response to increasing number of calls per individual; ‘i’ – response to increasing number of individuals; ‘y’- yes; ‘n’ – no; ‘+’ – increase; ‘-‘ – decrease; ‘ns’ – not significant, does not change with a parameter.

### Univariate metrics

Univariate metrics: F, PIC variants (PIC_betweentot_, PIC_betweenmeans_), H_S_ variants (H_Sntot_, H_Snpergroup_, H_Sngroups_, H_Svarcomp_).

All explored univariate metrics increased with increasing individuality in the data. However, only PIC_betweentot_, PIC_betweenmeans_, H_Snpergroup_ and H_Svarcomp_ estimates were independent of the number of calls and the number of individuals used to calculate the metric (Figure 3). These general patterns were qualitatively identical when all results were pooled or if only one of the parameters (number of calls, number of individuals, individuality) was changed at a time and the others were kept constant at the middle value (see Supplement 3 for detailed results including ANOVA tests).

All four sampling-independent metrics (PIC_betweentot_, PIC_betweenmeans_, H_Snpergroup_ and H_Svarcomp_) were highly correlated (Spearman correlation, all r > 0.99). H_Snpergroup_ and H_Svarcomp_ correctly converged to 0 in the case when individuality was set to be negligible (id = 0.01), while PIC_betweentot_ and PIC_betweenmeans_ converged to higher values (1.01 and 0.32 respectively). H_Svarcomp_ was equal to 2 * H_Snpergroup_ (see Supplement 4 for details). We further considered only the H_Snpergroup_ in multivariate analyses.

Overall, H_S_ performed best and best matched the characteristics of an ideal metric (Table 1).

**Figure 3.**
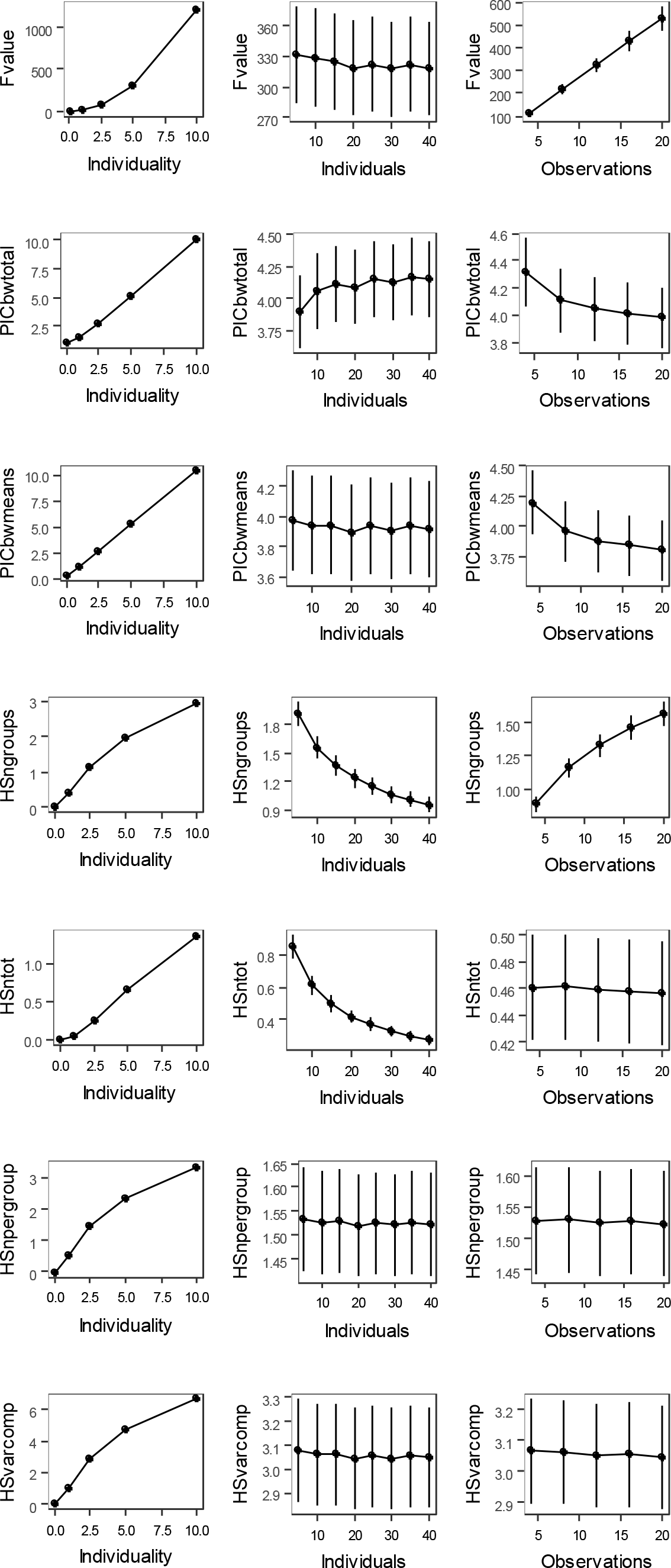
Variation in univariate identity metrics in response to artificial dataset parameters: individuality, number of calls per individual, and number of individuals. Means and 95% confidence intervals are shown. Graphs were plotted using all data pooled together.

### Multivariate metrics

The performance of multivariate identity metrics is illustrated in Figure 4. All metrics increased with increasing individuality. DS, H_S_, and MI increased with increasing number of variables available and decreased with increasing covariance between variables. Only H_M_ did not change in response to increasing the number of individuals. H_S_ and H_M_ did not change in response to increasing the number of calls per individual. These general patterns were qualitatively identical when all results were pooled or if one parameter was changed at a time and others were kept constant at the middle value (see Supplement 5 for detailed results including ANOVA tests).

**Figure 4.**
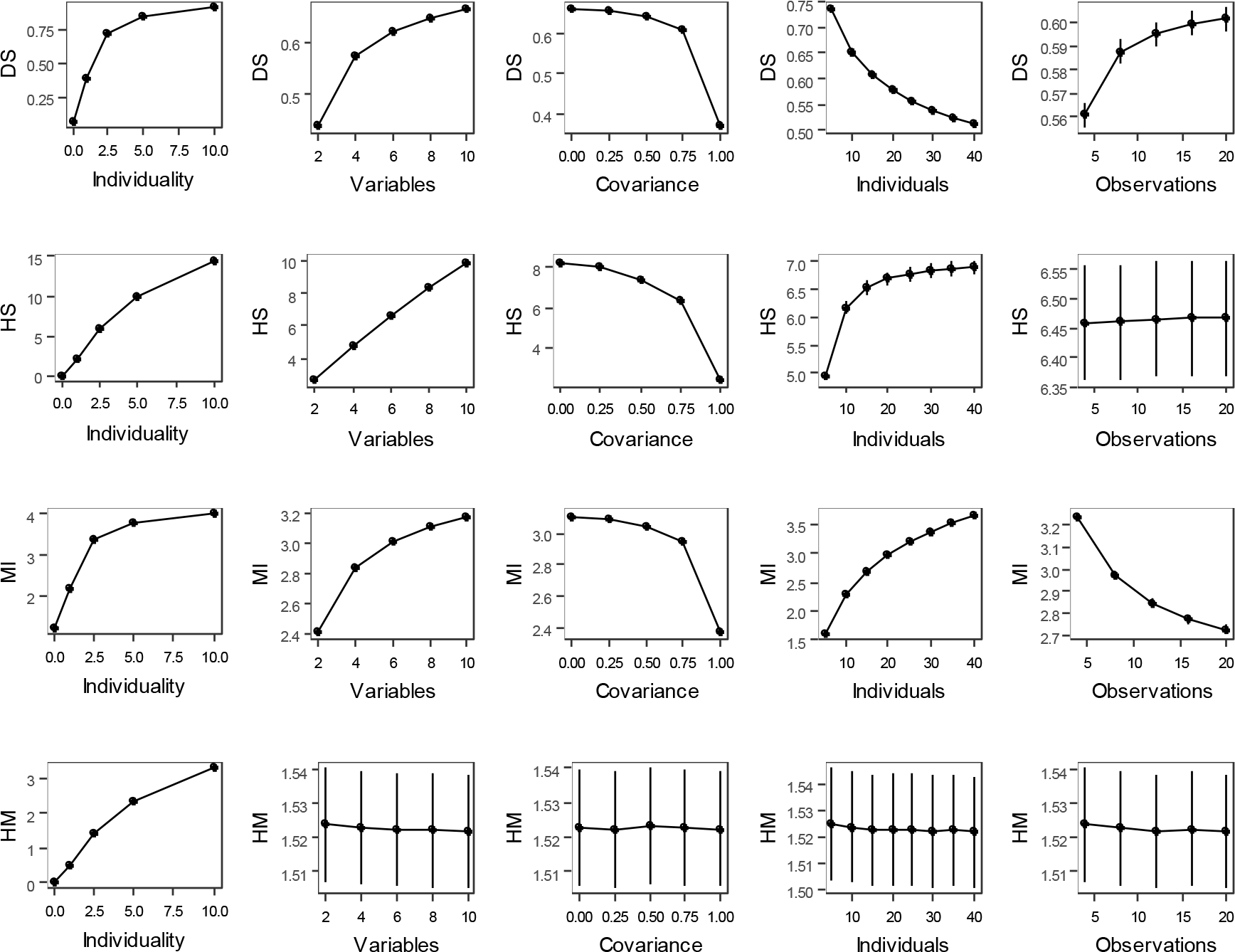
Multivariate identity metrics in response to changing individuality, covariance between variables, number of variables, number of calls per individual, and number of individuals in artificial data. Means and 95% confidence intervals are shown.

Despite the different response of metrics to some of the simulated parameters, there was still moderate to high agreement among metrics about identity content in the data (Spearman correlations, mean r ± SD = 0.82 ± 0.07; minimum r = 0.71 for correlation between DS and MI; maximum r = 0.95 for correlation between DS and H_S_). H_S_ had the greatest correlations with other metrics (average R = 0.88). We found no advantage to using H_M_ over H_S_ as previously suggested. Instead, H_M_ was equal to H_S_ per variable (H_M_ = H_S_ / p) (Supplement 6).

Thus, our simulations show that H_S_ performed best and matched the characteristics of the ideal metric in 6/7 cases, followed by H_M_ (5/7), DS (4/7), and MI (both 3/7) (Table 1).

### Potential for removing bias in H_S_

We observed no significant association between H_S_ and the number of individuals in the univariate case so the question arose about the precise cause of the bias in the multivariate case. This bias was only present when data were subjected to Principle Components Analysis (PCA). However, PCA is required to create uncorrelated components for H_S_ calculation. It is possible that the more variables measured, the more individuals need to be sampled in order to reduce this bias. We therefore fixed the number of variables to 5, 10, and 20 (p = 5, 10, 20) and varied the ratio of number of individuals to number of variables ‘i to p ratio’ from 0.5 to 5 (‘i to p ratio’ = 0.5, 1, 1.5, 2, 3, 5) by using different numbers of individuals in our simulations (i = 3, 5, 8, 10, 15, 20, 25, 30, 40, 50, 60, 100 depending on number of variables and “i to p ratio”). The number of calls per individual was set to 10. Individuality and covariance were both chosen randomly in each iteration from predefined intervals used in the earlier simulations (covariance range = [0, 0.25, 0.5, 0.75, 1]; individuality range = [0.01, 1, 2.5, 5, 10]). We used 100 and 1000 iterations for each ‘i to p ratio’ to get less and more conservative estimates. H_S_ did not rise significantly after the number of individuals reached at least the number of parameters in case of 100 iterations (One-way ANOVA F_5, 1794_ = 7.68, P < 0.001; no significant differences between levels if ‘i to p’ ≥ 1, all p > 0.132) (Figure 5), or at least twice the number of parameters in case of 1000 iterations (one-way ANOVA F_5, 17994_ = 63.19, P < 0.001; no significant differences between levels if ‘i to p’ ≥ 2, all p > 0.104).

**Figure 5.**
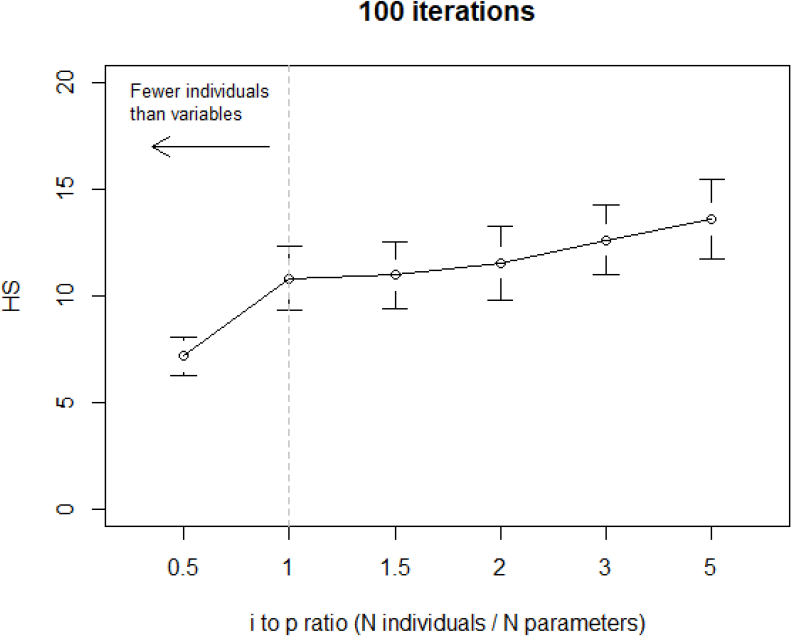
H_S_ and “i to p ratio” (number of individuals / number of variables) for situation with 100 iterations. H_S_ was under-estimated if there are fewer individuals than variables. Means and 95% confidence intervals are shown.

### Converting DS to H_S_ and vice versa

We used simple linear regression and non-parametric loess regression to estimate H_S_ based on DS and vice versa. There was a previously suggested linear relationship that had a limit of H_S_ = 8 where the DS values were 100% correct discrimination (Beecher 1989). Because the H_S_ values in our original simulated datasets far exceeded 8 in many cases (maximum H_S_ = 32.9), we generated a new set of simulated datasets with individuality ranging between 0.1 and 2 (id = 0.1, 0.25, 0.5, 0.75, 1, 1.33, 1.66, 2), covariance set to zero (cov = 0), number of iterations was reduced to 10 (it = 10), and other parameters were set as in previous models (p = 2, 4, 6, 8, 10; i = 5, 10, 15, 20, 25, 30, 35, 40; o = 4, 8, 12, 16, 20). These settings led to H_S_ values up to 13.0 for data used for model building, and H_S_ values up to 14.4 in the case of data used for model testing. These values are much closer to 8 and also much closer to H_S_ values reported from nature.

Loess models took into account specific sampling of the dataset; specifically, we included as predictors the number of calls per individual and the number of individuals. We compared the loess conversion and linear conversion models of DS and H_S_. In general, loess estimates were closer to the ideal prediction (intercept = 0, beta = 1) and the loess model reduced error of both DS and H_S_ estimates to about a half compared to linear estimates (Figure 6). Both H_S_ estimates were underestimated for high values of H_S_. The ceiling value is clearly apparent for linear estimates of H_S_. It is still visible in case of loess estimates but loess predictions remain reasonably good up to about H_S_ = 10.

**Figure 6.**
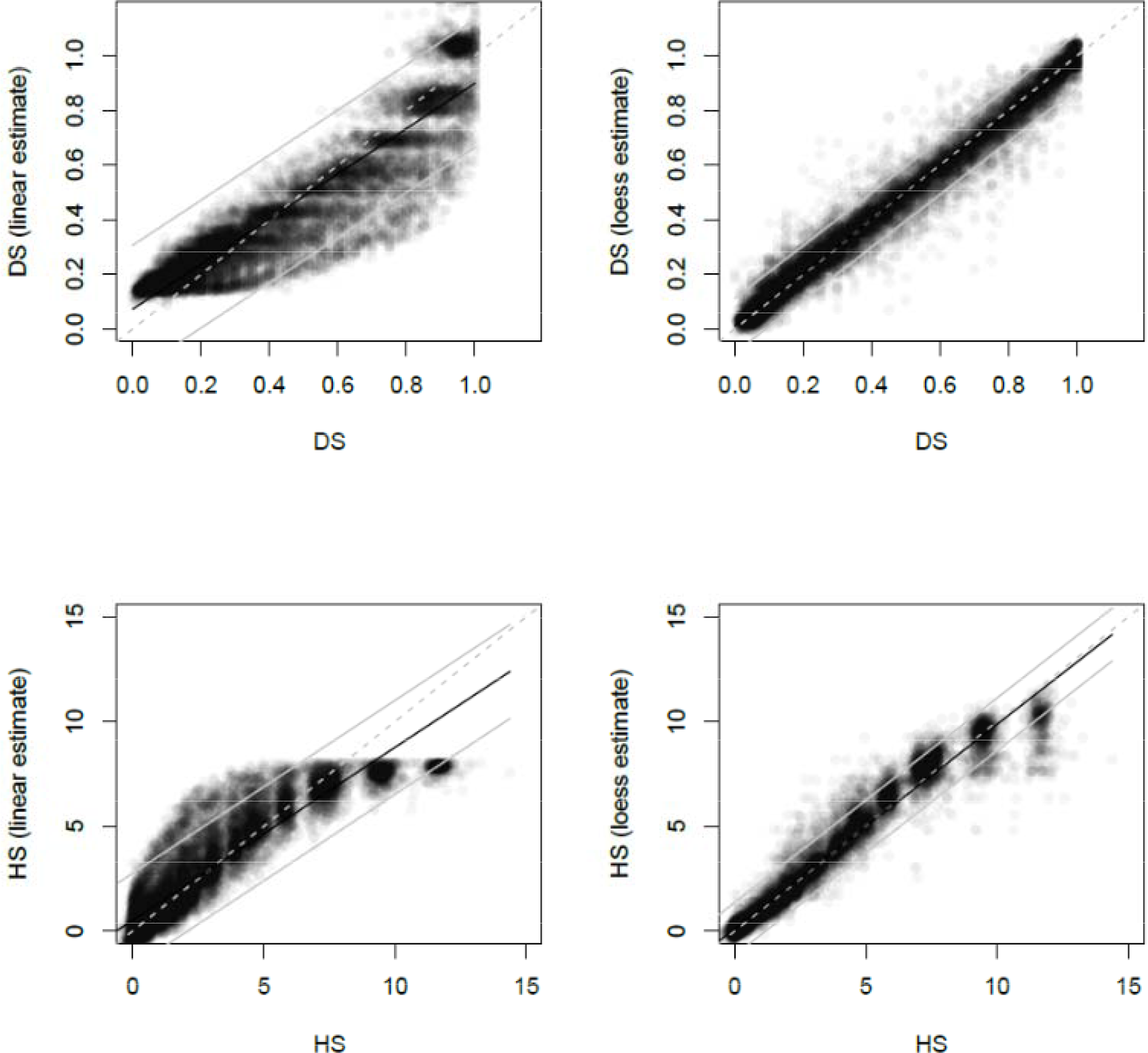
Estimation of H_S_ and DS based on linear and loess transformation of DS and H_S_ respectively for datasets with H_S_ up to 14.4. **Linear DS estimation**: Intercept = 0.07, Beta = 0,83, R^2^ = 0.83, Standard Error of Estimate (SEE) = 0.12, 95% Prediction interval = predicted value ± 0.23; **DS loess estimation**: Intercept = 0.01, Beta = 0.98, R^2^ = 0.97, Standard Error of Estimate (SEE) = 0.05, 95% Prediction interval = predicted value ± 0.10. **Linear H_S_ estimation**: Intercept = 0.51, Beta = 0.83, R^2^ = 0.83, Standard Error of Estimate (SEE) = 1.14, 95% Prediction interval = predicted value ± 2.24; **HS loess estimation**: Intercept = 0.11, Beta = 0.98, R^2^ = 0.95, Standard Error of Estimate (SEE) = 0.64, 95% Prediction interval = predicted value ± 1.26.

### Correlations between calculated and estimated metrics

We were further interested in how H_Sest_ and DS_est_ might represent H_S_ and DS of a particular sample of individuals or H_Sfull_ and DS_full_ of the whole population. For this purpose, we first generated 50 full datasets with different identity levels representing 50 hypothetical populations of different species. Each dataset comprised of 40 individuals, 20 calls per individual, and 10 parameters. For these datasets, individuality was set randomly ranging between 0.2 – 2 (0.1 increments), and the covariance was set randomly ranging between 0.2 – 0.8 (0.1 increments). These settings generated datasets with H_Sfull_ values that ranged from 0.22 – 9.89 (mean ± sd: 4.72 ± 2.95). Then, we repeatedly subsampled these datasets to get partial datasets which simulate different sampling of the population. We subsampled 5-40 individuals and 4-20 calls per individual per dataset in each of total 20 iterations. We also repeatedly subsampled our empirical datasets. We subsampled 5-33 individuals and 4-10 calls per individual per dataset in each of total 20 iterations. The number of parameters was not randomized – we always kept the original number of variables.

In simulated datasets, H_S_ and H_Sest_ were correlated almost perfectly with each other and with H_Sfull_ (all average Pearson r > 0.97). There was no difference among correlation coefficients from correlations between H_Sfull_, H_S_, and H_Sest_ (Friedman Chi Square = 3.6, p = 0.165). In empirical datasets, H_S_ calculated on partial datasets still reflected the H_Sfull_ almost perfectly (average Pearson r = 0.99). While H_Sest_ reflected H_S_ of partial dataset (average Pearson r = 0.90), and H_Sfull_ (average Pearson r = 0.88) was slightly worse, it remained a reasonable fit. However, H_Sest_ did not reflect H_Sfull_ as precisely as it did H_S_ (Friedman Chi Square = 33.6, p < 0.001, post-hoc test: H_S_ - H_Sfull_ vs. H_Sest_ - H_Sfull_, p < 0.001).

DS in simulated datasets, was almost perfectly correlated with DS_est_ (average Pearson r = 0.99). Although the relationship between DS and DS_est_ was significantly worse in a full dataset (DS_full_) (Friedman Chi Square = 40.0, p < 0.001; both post-hoc tests: p < 0.005), these associations remained strong (DS_full_ and DS: average Pearson r = 0.95; DS_full_ and DS_est_: average Pearson r = 0.96). In empirical datasets, the correlation between DS and DS_est_ was lower than in case of artificial datasets (average Pearson r = 0.91). DS and DS_est_ of partial datasets had comparable correlations to DS_full_ (DS_full_ and DS: average Pearson r = 0.88; DS_full_ and DS_est_: average Pearson r = 0.86). Thus, the performance of DS and DS_est_ to reflect each other or DS_full_ did not differ (Friedman Chi Squre = 0.9, p = 0.638).

## Discussion

All identity metrics had systematic biases that emerged from sampling decisions. Biases induced by the number of individuals and the number of calls per individual in a sample both decreased with improving sampling. H_S_ was closest to an ideal identity metric in the univariate case when identity was assessed for a single variable, as well as in multivariate case when identity was assessed for a set of several different variables. The bias caused by the number of individuals in the sample used to calculate H_S_ could be removed by having at least the same number of individuals as the number of variables. H_S_ was the most consistent metric and best correlated with DS and other identity metrics. H_S_ could be converted reliably into DS and vice-versa.

### Univariate identity metrics

Beecher’s information statistic (H_S_) (Beecher et al., 1986; Beecher, 1989) and Potential for individual coding (PIC) (Robisson et al., 1993; Lengagne et al., 1997) were both suggested as unbiased alternative metrics to F values. We confirmed that both H_S_ (when calculated properly) and PIC provide unbiased estimates of identity information. Further, we show that these two metrics are almost perfectly correlated and, hence, in general, they both measure the same thing. PIC reflects the number of potential individual signatures within a population in same way as 2^*H*_*S*_^ does. However, PIC slightly differs from H_S_ and deviates from expected zero values if there is low identity content in a signal that approaches zero. It is important to realize that variables with PIC_betweentot_ value > 1 need not convey meaningful individual information as commonly assumed. Using the PIC_betweentot_ does not create overly spurious conclusions but rather including more less-important variables increases noise in subsequent analyses. Studies using the number of individuals as ‘n’ to calculate H_S_ most likely under-estimates the real H_S_ value because the number of individuals is typically higher than the number of calls per individual in those studies. H_S_ has been suggested as a suitable metric for comparative analyses and H_S_ has been used for such purposes in a few such analyses. We think the overall conclusions of these analyses are valid whenever the same sampling protocol was used across species (e.g., Pollard & Blumstein, 2011).

### Multivariate identity metrics

Discrimination score (DS) is by far the most used acoustic identity metric, despite numerous studies showing systematic biases in DS (e.g., Beecher, 1989; Bee, Kozich, Blackwell, & Gerhardt, 2001; Budka, Wojas, & Osiejuk, 2015; Linhart & Šálek, 2017). We conclude that Beecher’s information statistic (H_S_) (Beecher, 1989) is the best of the several alternative metrics proposed. In addition to H_S_, two other metrics – H_M_ and MI – were introduced to overcome biases of discrimination scores. We did not find that H_M_ or MI were better suited than H_S_. Unfortunately, performance of neither of H_M_ or MI was directly compared, nor was either shown to exceed the performance of H_S_ (Searby & Jouventin, 2004; Mathevon et al., 2010) despite the fact that both are grounded in information theory and use the same measurement unit (bits) as H_S_. The robustness of H_M_ towards sampling reported here (number of individuals, number of calls, even number of variables and covariance) could be seen as attractive. However, as we show, H_M_ quantifies identity information per variable and not the identity information of the entire signal. If one is interested in total identity information, with H_M_, it is necessary to know the effective number of variables (i.e., if there is perfect covariance between the variables, the effective number of variables is 1 no matter how many variables are used), which can be difficult in real situations. Mutual information (MI) is derived from confusion matrix of discrimination analysis and we show it has similar shortcomings as discrimination scores. Our results showing biases in MI are in line with previous studies that investigated measures of clustering for various machine learning purposes where potentially unbiased variants of MI are searched for (Marrelec, Messé, & Bellec, 2015; Amelio & Pizzuti, 2017).

Although we suggest that H_S_ should be generally used to quantify individuality, some questions on identity signaling might still need to rely on the other identity metrics or approaches. For example, researchers might be interested in whether distinctiveness of individuals increases during ontogeny (Briefer & McElligott 2012, Lapshina et al., 2012, Syrová et al., 2017). In such cases, assessment on individual level is required (distances, discrimination score) while H_S_ would only provide overall identity information for each ontogeny stage making further statistical assessment impossible.

### Precision of conversion between metrics

Both H_S_ and H_M_ values were previously found to correlate well with DS (Beecher, 1989; Searby & Jouventin, 2004). We extend these previous findings on H_S_ (Beecher, 1989) to situations with unequal sampling and we show it is possible to convert between H_S_ and DS with an acceptable amount of error even when datasets differ in the number of individuals and calls per individual. Predicting DS from H_S_ has an advantage of being more precise than predicting H_S_ from DS. The precision of conversion decreased in real datasets compared to simulated datasets. However, the decrease was not dramatic, especially when considering that the conversion model was derived from simulated datasets with only two uncorrelated variables while real datasets differed in both the number of variables and their covariance structure. Furthermore, real datasets had issues associated with multivariate normality, which is a common problem of many studies and which also likely worsened the conversion precision and metric consistency.

### Identity metrics in comparative analyses

Despite the systematic biases related to sample size in DS (the most often used metric) and in H_S_ (the best metric), we show that these biases, while introducing certain level of noise, may not be fatal to those who desire to compare identity between individuals or species because our H_S_ and DS values based on an entire population or subsamples from these populations were well correlated for both simulated and empirical datasets.

### Sample size considerations

Biases of both DS and H_S_ decrease with increasing sample sizes. Researchers using DS as an identity metric have been warned about the problems with low sample sizes. However, these concerns were generally related to the number of observations per group (typically, calls per individual) (Mundry & Sommer, 2007). Indeed, it has also been frequently pointed out that PCA is sensitive to sample sizes. However, the sample size recommendations typically relate to the total sample size (e.g., McGarigal, Cushman, & Stafford, 2000), while applying PCA to identity research is somewhat special and assumes that principle components reflect the variation between individuals. Our study suggests that number of individuals should always be at least as large as number of variables whenever PCA is used to study individual identity.

### Using identity metrics across modalities

We evaluated the efficacy of all metrics within the acoustic modality only. It is increasingly recognized that signals may employ multiple modalities (Partan & Marler, 1999; Proops, McComb, & Reby, 2009; Pitcher, Briefer, Baciadonna, & McElligott, 2017). There is no reason to believe that modality constrains the use of these metrics and, in principle, all of the identity metrics could be used in visual or chemical domains as well (Beecher, 1982; Beecher, 1989; Kondo & Izawa, 2014). However, identity information outside the acoustic domain is rarely quantified with the metrics described here because they all require assessment of a signal’s within individual variation. The reasons might be that other modalities are assumed to be more static or because of technical difficulties in quantifying within-individual variation. The latter seems to be a case. The latest progress in machine learning and image analysis suggests that it should be possible to conduct individual discrimination tasks in a similar way to that used for acoustic signals (Allen & Higham, 2015; Van Belleghem et al., 2018). Finally, repeated sampling of individual signatures in olfactory secretions is becoming more common (Kean et al., 2015; Deshpande, Furton, & Mills, 2018). Thus, researchers may try to quantify potential individual identity information in visual and chemical signals in future studies.

### Conclusion

We have shown that H_S_ is the identity metric with the best performance in both univariate and multivariate contexts. Given that H_S_ may not be sufficient in all cases, we encourage further research to develop new metrics to quantify identity information in signals. However, new metrics should always be appropriately assessed and their performance directly compared to the best existing metrics. The datasets and algorithms we have provided should aid in future comparisons.

## Supporting information

Supplement 6 - HS and HM relationship

Supplement 5 - Multivariate Metrics-data pooled or by single parameter changing

Supplement 4 - Relationship between HS and PIC

Supplement 3 - Univariate Metrics-data pooled or by single parameter changing

Supplement 2 - Univariate and multivariate simulated datasets

Supplement 1 - Overview of individual identity metrics

## Acknowledgements

PL received funding from the European Union’s Horizon 2020 research and innovation programme under the Marie Skłodowska-Curie grant agreement No. 665778 administered by the National Science Centre, Poland (UMO-2015/19/P/NZ8/02507). DTB is supported by the NSF. MŠp, MS, and RP were supported by Czech Science Foundation (GA14-27925S) and Czech Ministry of Agriculture (MZE-RO0718). MŠá work was supported by the research aim of the Czech Academy of Sciences (RVO 68081766).

## Data Accessibility statement

Data and code used for this article are available at GitHub and ZENODO public repositories under permissive free software MIT license (Linhart 2018).

