## Supplement 6 - HS and HM relationship for "Measuring individual identity information in animal signals: Overview and performance of available identity metrics"

### The relationship between H_S_ and H_M_

Searby and Jouventin (2004) suggested that H_M_ should be favored over H_S_ in case of modulated signals. However, they only presented the relationship between H_M_ and discrimination score and did not directly compare the performance of H_S_ and H_M_. Moreover, their artificial dataset was limited in the number of manipulated parameters. We were therefore interested in better understanding the relationship between the H_M_ and H_S_ in our data and whether H_M_ could have advantages over H_S_.

We first tried to replicate results of Searby and Jouventin (2004) but included the calculation of H_S_. The process and functions necessary to replicate the data are described in the file “replicate Searby Jouventin 2004.R” within the provided R project (Linhart, 2018). We were able to replicate the results of Searby and Jouventin (2004) quite well (Figure S6.1a). H_S_ was almost identical to H_M_ * 25 (Figure S6.1b). Because the dataset comprised 25 identity variables, H_M_ could be potentially seen as the average H_S_ per identity variable. We further tested this possibility in our datasets (see below). We found no advantage of using H_M_ over H_S_ for modulated signals as the two metrics are perfectly correlated in the Searby and Jouventin (2004) replicated dataset (R^2^ = 1.00, Figure S6.1b).

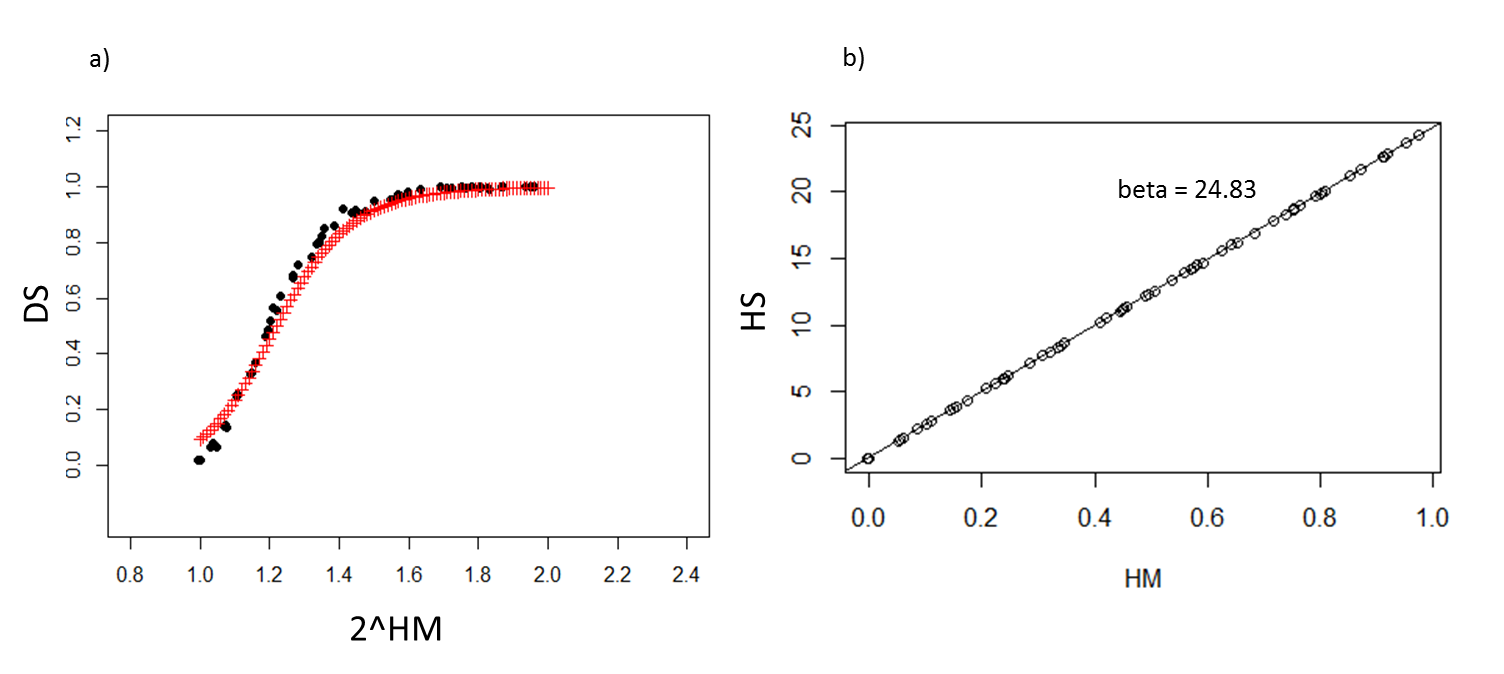

**Figure S6.1**. Relationship between discrimination score (DS) and 2^H_M_ as reported by Searby and Jouventin (2004) and our replication of the same data (a) – red crosses show the line predicted by Searby and Jouventin (2004, Fig 2b). Relationship between H_S_ and H_M_ in replication of data from Searby and Jouventin 2004 (b).

However, in our dataset, this perfect relationship can became blurred because of changes in covariance (H_S_ values are lower than expected from H_M_) and insufficient sampling (in cases when there are few individuals, H_S_ is underestimated) (Figure S6.2a). When the covariance is set to zero and with sufficient sampling of individuals (full set of 40 individuals) H_S_ is again almost equivalent to H_M_ * number of variables (Figure S6.2b). We suggest that H_S_ should be preferred over H_M_ because H_M_ requires additional information about the dimensionality of the data which would need to be assessed separately for non-independent variables.

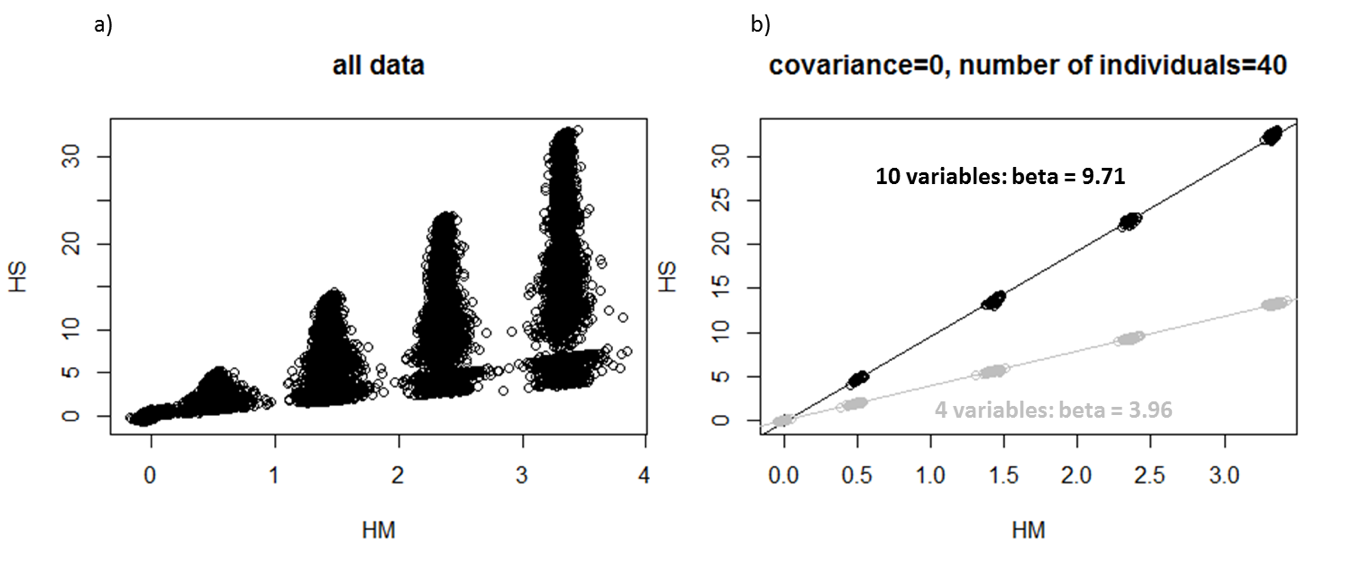

**Figure S6.2.** Relationship between H_M_ and H_S_ in our all simulated datasets for all data pooled (a) and for selected datasets with covariance=0 and number of individuals=40 (b).
