## Supplement 4 - Relationship between HS and PIC for "Measuring individual identity information in animal signals: Overview and performance of available identity metrics"

### Relationship between H_S_ variants and PIC variants

PIC_betweentot_, PIC_betweenmeans_, and 2^ H_Snpergroup_ are almost identical metrics as revealed by high linear correlations between them (R^2^(PIC_betweentot_ vs. PIC_betweenmeans_) = 0.996; R^2^(PIC_betweenmeans_ vs. 2^H_Snpergroup_) = 0.997; R^2^(PIC_betweentot_ vs. 2^H_Snpergroup_) = 0.997). However, they differ in small important details. If there is essentially no individual variability in the data (individuality = 0.01), H_Snpergroup_ correctly gives values around 0 (mean ± sd = 0.00 ± 0.05) while both PICs converge to higher values (PIC_betweentot_ mean ± sd = 1.01 ± 0.05; PIC_betweenmeans_ mean ± sd = 0.32 ± 0.13) (see Figure S4.1 below).

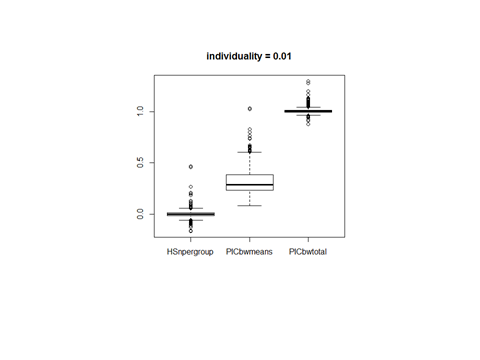

Metric Value

H_Snpergroup_	PIC_betweenmeans_	PIC_betweentot_

**Figure S4.1.** Values of H_Snpergroup_, PIC_betweenmeans_, and PIC_betweentot_ for 0 individuality in data.

H_Snpergroup_ and H_Svarcomp_ were both independent of sampling but their absolute values were different. Interestingly, H_Svarcomp_ was almost identical to 2 * H_Snpergroup_ (linear regression: R2 = 1, intercept = 0.01, beta = 1.99).
