## Supplement 2 - Univariate and multivariate simulated datasets for "Measuring individual identity information in animal signals: Overview and performance of available identity metrics"

### Detail description of preparation of simulated datasets

**Univariate simulated datasets.** First, we generated individual means for a predefined number of individuals “i” (normal distribution, “rnorm” function, mean = 1000, SD_between_ = 1). We manipulated identity information in the dataset by changing the within individual variation (SD_within_) and we generated a predefined number of random observations “o” around each individual mean (normal distribution, “rnorm” function, mean = individual mean, SD_within_ = SD_between_ / individuality “id”). The number of individuals “i” ranged from 5 to 40 (i = 5, 10, 15, 20, 25, 30, 35, 40), number of observations “o” ranged from 4 to 20 (o = 4, 8, 12, 16, 20), and individuality “id” ranged from 0.01 for no individuality to 10 for very high individuality (id = 0.01, 1, 2.5, 5, 10).

**Multivariate simulated datasets.** First, we generated a matrix representing mean individual values of variables for each of the individuals (multivariate normal distribution, “mvrnorm” function, mean for each variable = 0, variance-covariance matrix). Variances on the diagonal of the covariance matrix were set equal to 1 (hence SD_between_ = 1) and all covariances between variable pairs were set equal to the predefined covariance “cov”. Then, we generated a predefined number of random observations “o” around each individual and a variable mean (“rnorm” function, mean = individual mean, SD_within_ = SD_between_ / individuality “id”). The number of individuals “i” ranged from 5 to 40 (i = 5, 10, 15, 20, 25, 30, 35, 40), number of observations “o” ranged from 4 to 20 (o = 4, 8, 12, 16, 20), and individuality “id” ranged from 0.01 for no individuality, to 10 for very high individuality (id = 0.01, 1, 2.5, 5, 10) as in case of univariate datasets (see Figure 1 for illustration of individuality). The number of variables “p” ranged from 2 to 10 (p = 2, 4, 6, 8, 10) and covariance “cov” ranged from 0 for complete independence of variables to 1 for absolute correlation between variables (cov = 0, 0.25, 0.5, 0.75, 1).

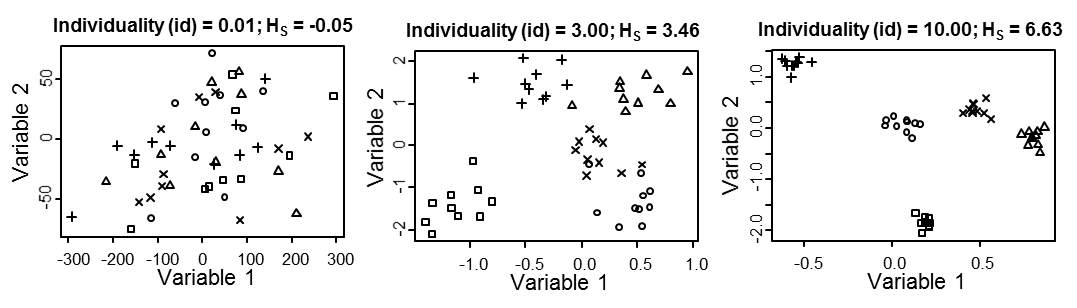

**Figure 1**. Illustration of three artificial multivariate datasets that differ only in the individuality used to generate datasets. Settings for the function generating these datasets: i = 5, o = 10, p = 2, cov = 0, id = 0.01, 3, and 10
