## Supplement 1 - Overview of individual identity metrics for "Measuring individual identity information in animal signals: Overview and performance of available identity metrics"

### F-value (F, univariate)

The earliest and the most simple way of testing for individual differences in signals is to fit a one-way ANOVA with individual as the independent variable to explain variation in some signal attribute (e.g., Miller, 1978). F is calculated by dividing between group mean squares MS(B) by within group mean squares MS(W):

$$F=\frac{MS(B)}{MS(W)}$$

(1)

The well recognized problem with F-values, however, is that F-values are systematically influenced by sampling; thus, they are not well suited for comparisons between studies. The two following metrics can overcome this shortcoming.

### Potential of individual coding (PIC, univariate)

Potential of individual coding (PIC) is defined as a ratio of between-individual to within-individual coefficients of variation (e.g., Robisson, Aubin, & Bremond, 1993; Lengagne et al., 1997):

$$PIC=\frac{{CV}_{b}}{{CV}_{w}}$$

(2)

Like the F-value, PIC is mainly used to compare the level of individual variation within each signal variable. For example, PIC was used to show that frequency variables are better suited to convey individuality than temporal variables in king penguins, *Aptenodytes patagonicus* (Robisson, 1992). However, PIC has some shortcomings. It can only handle variables on a ratio scale (Robisson et al., 1993). Therefore, it cannot handle variables that can have negative and positive values like, for example, the slope of a frequency modulation. For this reason, PIC also cannot be used with uncorrelated variable components of original data obtained from Principal component analysis. Furthermore, it seems that researchers have used two slightly different ways to calculate PIC. CV_b_ in the formula is either calculated from all values (PIC_betweentot_) ( e.g., Owens & Freeberg, 2007; Charrier et al., 2010; Reers, Jacot, & Forstmeier, 2011; Pettitt, Bourne, & Bee, 2013; Salmi, Hammerschmidt, & Doran-Sheehy, 2014), or means for each individual are calculated first and CV_b_ is then calculated from these means (PIC_betweenmeans_) (e.g., Lingle, Rendall, & Pellis, 2007; Lein, 2008; Reichert, 2013).

### Beecher’s statistic (H_S_, univariate / multivariate)

H_S_ is grounded in information theory (Beecher, 1989) and it is measured in bits. H_S_ is based on F-value but unlike F-value, H_S_ accounts for sample size:

$$H_{S}={log}_{2}\sqrt{\frac{F+n-1}{n}}$$

(3)

It can be used for a single variable, or multiple variables can be subjected to Principal components analysis (PCA), H_S_ calculated and summed over each principal component of an entire multivariate signal. A strength of H_S_ is that it permits the estimation of a number of possible unique individual signatures within a population. Different variants of this equation have been used in literature. The source of confusion is the ’n’ in the formula. Total sample size (H_Sntot_), number of groups (i.e., individuals) (H_Sngroups_), and number of samples per group (H_Snpergroup_) could all be used as ‘n’ in this equation. Some studies explicitly state they used number of individuals as ‘n’ (e.g., Pollard, Blumstein, & Griffin, 2010; Linhart & Šálek, 2017), but the properties of H_S_ values in these studies did not match the properties suggested in the original article by Beecher (1989). Yet another approach to calculate H_S_ is to extract the variance component estimates and use the total (ơ_T_) and the residual variance (ơ_W_, associated with random factor) to calculate H_S_ (H_Svarcomp_) (Beecher, 1989; Carter, Logsdon, Arnold, Menchaca, & Medellin, 2012):

$$H_{S}={log}_{2} \frac{\sigma_{T}}{\sigma_{W}}$$

(4)

### H for modulated signals (H_M_, multivariate)

It has been argued that H_S_ is good for scalar variables and is not suitable in case of modulated signals (Searby & Jouventin, 2004). Therefore, H_M_ was introduced to quantify the individuality of modulated signals. H_M_ is derived from H_S_. Unlike H_S_, H_M_ does not consider each measured trait (or principal component) but uses multivariate Euclidean distances between samples (dist_T_ = sum of distances of all samples from their centroid; dist_W_ = sum of distances of samples within individual to its centroid) to estimate F in the original formula:

$$H_{M}={log}_{2}\sqrt{\frac{F_{M}+n-1}{n}}$$

(5)

$$F_{M}= \frac{n-1}{g-1}* \left( \frac{{dist}_{T}-g*{dist}_{W}}{{dist}_{W}} \right)$$

(6)

where ‘n’ = number of observations and ‘g’ = number of groups (individuals). H_M_ has been rarely used, and while H_M_ values reflect the possibility of discrimination by humans and discriminant analysis, its superiority and relationship to H_S_ was not directly shown.

### Discrimination score (DS, multivariate)

To quantify individuality in acoustic signals, most researchers have used discriminant function analysis and calculated discrimination scores (e.g., Hafner et al., 1979). Typically, different features of a signal are measured. These variables (raw or after various transformations) are then subjected to discriminant analysis and the total percentage of calls assigned to the correct individual is reported. Several disadvantages of discrimination scores were previously reported including the undesirable effect that increasing the number of individuals in a sample systematically decreases discrimination. By contrast, more calls per individual can improve discrimination scores. Due to these biases, researchers were reluctant to carry out quantitative comparisons among different studies (Insley et al., 2003). Despite reported biases, discrimination scores are widely used, probably because of the intuitive association with the task of individual recognition.

### Mutual information (MI, multivariate)

Mutual information (MI) may represent an intermediate step between DS and H_S_ and DS and H_M_. Similar to H_S_ and H_M_, mutual information is grounded in information theory and expressed in bits. However, it is calculated from a confusion matrix and has been suggested as a metric of goodness of classification that is independent of the number of calls and number of individuals in the sample (Mathevon et al., 2010).

$$MI= \sum_{i,j} {log}_{2}\left( \frac{p(i,j)}{p\left( i \right)\cdot p(j)} \right)$$

(7)
